# Novel introductions of human-origin H3N2 Influenza viruses in Swine, Chile

**DOI:** 10.1101/2024.05.08.593172

**Authors:** B. Agüero, N. Ariyama, Felipe Berrios, Nikita Enciso, Barbara Quezada, R.A. Medina, V. Neira

## Abstract

Influenza A virus (IAV) continuously threatens animal and public health globally, with swine serving as a crucial reservoir for viral reassortment and evolution. In Chile, H1N2 and H3N2 subtypes were introduced in the swine population before the H1N1 2009 pandemic, and the H1N1 was introduced from the H1N1pdm09 by successive reverse zoonotic events. Here, we report two novel introductions of IAV H3N2 human-origin in Chilean swine during 2023. Our study reveals a closer relationship between recent human seasonal H3N2 and novel swine strains. Interestingly, one strain maintains all the genes from the original human virus, but the other strain is already a reassortment of human H3N2 and an H1N2 previously observed on the farm. Observing global IAV sequences, a similar pattern was identified in the USA confirming the reverse zoonotic potential of current seasonal human H3N2 strains. These results highlight the importance of ongoing surveillance and reinforce biosecurity in swine farms. These findings raise questions about their potential impact on viral dynamics in the swine population and public health, underscoring the need for further investigation into the origin and evolutionary dynamics of this emerging swine H3N2 reassortant virus.

## 1 Introduction

Influenza A virus (IAV) is swine an important concern for animal health and also for zoonotic potential. Swine, susceptible to avian and human strains, is pivotal as a “mixing vessel” for influenza virus reassortment and evolution (1). Swine is highly susceptible to being affected by human strains (reverse zoonotic), and exhibits high genetic and antigenic diversity of IAV, facilitated by reassortment and antigenic drift, which pose challenges for vaccine efficacy and zoonotic risk assessment (2). This can lead to the emergence of new strains or subtypes with zoonotic potential, as happened in 2009 with the appearance of the novel reassortant H1N1 influenza virus, which had genes from porcine, avian, and human origins (3–5). Economically, swine influenza outbreaks can also generate significant economic losses for pig production by decreased productivity, and increased healthcare costs(6), highlighting the ongoing need for effective disease management strategies to safeguard animal welfare and economic stability.

In Chile, the presence of swine influenza subtypes H1N1, H1N2, and H3N2 has been documented in intensive production facilities (7,8), as revealed by an extensive study spanning over a decade. We collected and analyzed more than 10,000 samples from swine populations across the country during this period. In detail, we have identified strains such as H1N1pdm09 and H1N2 clade 1.B, as well as H3N2. The H1N1pdm09 strains were recently introduced from the human population after its pandemic, and H1N2s and H3N2s were also transmitted from humans to swine in various instances between 1980 and 2006 (seasonal human strains)(9). Remarkably, our datasets do not record any recent introductions of H3N2 viruses of human origin until now.

This study aims to characterize two newly identified human-origin H3N2 viruses found in Chile’s swine-intensive populations. The viruses were isolated from samples collected from intensively farmed pigs in Chile in 2023. This discovery underscores the ever-evolving nature of influenza viruses and emphasizes the vital need for ongoing surveillance and research efforts.

## 2 Materials and methods

### 2.1 Sample collection

To gain a comprehensive understanding of this study, we included all Influenza A Virus (IAV) samples collected from January 2023 to January 2024. These samples were predominantly collected by veterinarians in response to suspected IAV cases, but also in scheduled surveillance programs. A total of 1195 samples from 36 farms, representing over 90% of the national swine-intensive production, were submitted for IAV detection. Samples analyzed included nasal swabs (NS), oral fluid (OF), tracheal swabs (TS), bronchial swabs (BS), and lung tissue (L). The samples were sent on the same day or overnight using ice packs and polystyrene boxes. The samples were processed at the Animal Virology Laboratory of the Faculty of Veterinary and Animal Sciences at the University of Chile, which is recognized by swine practitioners as a national reference for influenza diagnosis. To maintain confidentiality, individual farms were not identified in this study.

### 2.2 Diagnostic and sequencing

The RNA extraction was performed using the Chomczynski-phenol solution (Winkler, BM-1755, Chile), followed by real-time RT-PCR targeting highly conserved regions of the IAV matrix (M) (10) using iTaq Universal Probes One-Step Kit (Bio-Rad, 1725141, USA) following the manufacturer’s instructions. Samples with a cycle threshold (Ct) value less than 35 were considered positive, and Ct values greater than 35 were considered negative. A subset of positive samples, with the lowest Ct values per case, were isolated and subtyped to detect H1, H3, N1, and N2 (11). The viral isolation was performed in Madin-Darby Canine Kidney (MDCK) cells (8,12). Two novel H3N2 suspected cases were attempted for whole genome sequencing using Oxford ONT technology at the Animal Virology Laboratory of the Faculty of Veterinary and Animal Sciences at the University of Chile.

Briefly, the IAV genome was amplified through a multi-segment one-step RT-PCR(13,14), and the libraries were prepared using the Native Barcoding Kit (SQK-NBD114.96), to be loaded in ONT Flonge flow cells (14). Reads were further assembled using the automated cloud-based pipeline CZ ID (15), by Genome Detective (16) and assembly using reference sequences in Geneious Prime® 2023.2.1.

### 2.3 Genetic Analysis

The genetic analysis was performed in all segments. The obtained sequences were queried in BLASTn (https://blast.ncbi.nlm.nih.gov/Blast.cgi) to identify the closest published sequences. The final data set for the genetic analysis included closely related sequences retrieved from BLAST, close sequences from Chilean-human, and all global human-origin swine H3N2 classified as human seasonal identified in OctoFLU (17). The reference sequences were obtained in public depositories GenBank and GISAID (18). Before the phylogeny, pairwise comparisons were performed between the novel H3N2 and reference sequences aligned using Clustal Omega (19). The phylogeny was inferred using Maximum likelihood by RAxML 8.2.11, using the GTR Gamma model with a bootstrap of 1,000 replicates in Geneious Prime® 2023.2.1. (20). The final trees were visualized in FigTree v1.4.4 (21).

In addition, to visualize reassortment events, we conducted recombination analysis by concatenating sequences. Briefly, the suspect reassortant strain was concatenated according to segment number, and potential parental strains were identified through BLAST analysis and consideration of whole genome availability. The analysis was executed using Dual Brothers (22) within the Geneious platform (20).

## 3 Results

A total of 1,195 samples were collected with an overall positivity rate of 10.6%. The most common samples were nasal swabs with 8.4% positivity, bronchial swabs with 35.9% positivity, and oral fluids with 37.5% positivity (Supplementary Table 1). Twenty out of the 36 farms analyzed were confirmed positive for IAV RT-PCR, obtaining 55.5% positivity. In general, the results were in concordance with the historical records of each farm, in terms of subtypes. However, in two cases we found evidence of H3N2, which is not common in Chile. These cases corresponded to two different farms not related epidemiologically. For case 1, this corresponded to a nursery site with endemic H1N1pmd09, the veterinarian mentioned an unexpected increase in clinical signs in 7-week-old pigs. This case was submitted in July 2023, including 9 nasal swabs and two lungs. The sample with the lowest Ct value was a nasal swab (Ct 24.2, gene M RT-PCR), and the subtyping was suspect for H3. Isolation was achieved on the second passage and was further sequenced, the characterized isolated was named A/swine/O’Higgins/VN1401-7442/2023 (H3N2).

The case 2, was also observed in nursery pigs. In this case, in October 2023. The submission included 4 nasal swabs, with 2 positive samples with low Ct values (13.7 and 15.5). The subtyping RT-PCR only detected the N2. The viral isolation was unsuccessful, so the virus was partially sequenced by Sanger using the direct sample identifying the subtype as H3. This farm was sampled again on January 25^th^, 2024, in this submission 4 out of 5 nasal swab samples were positive with Ct values ranging from 20.8 to 31.8. Similarly to the previous sampling, the samples were partially subtyped. In this submission, we were able to isolate the virus which was further sequenced. The characterized isolated was named A/swine/O’Higgins/VN1401-7826/2024 (H3N2).

Both isolates were successfully sequenced. For case 1, for the isolate A/swine/O’Higgins/VN1401-7442/2023 (H3N2), the blast results indicated that the closest sequences were viruses collected from humans in the United States during 2022 with an identity of 99.3% for the HA gene and higher for the rest. Also, several sequences collected from Chilean humans presented high identity which were included in the final tree. In contrast, for the A/swine/O’Higgins/VN1401-7826/2024 (H3N2) only the HA gene had the closest sequence from humans in the United States during 2022, with 98.98% identity, while the remaining genes were closely related to H1N2s viruses from the same company collected in previous years (See Supplementary Table 2). The pairwise comparison for H3 between the novel Chilean H3N2 and the 6 H3N2 isolates collected from Chilean swine in earlier years (9) had a low identity with the novel H3, with percentages < 88%.

The final trees for HA phylogeny were built with 62 sequences. For the HA, the novel H3N2 Chilean sequences belonged to a subcluster containing H3N2 isolates collected recently from humans and swine. All those sequences corresponded to human seasonal H3N2 collected in 2022, most of which corresponded to human sequences from the USA and Chile (Figure 1). For the NA, the strain A/swine/O’Higgins/VN1401-7442/2023 (H3N2) had the same pattern as the HA tree, which from a cluster with human sequences from the USA and Chile, corresponded to human seasonal H3N2 collected in 2022. In contrast, the A/swine/O’Higgins/VN1401-7826/2024 (H3N2) (Case 2), clustered with swine isolates of the H1N2 subtype collected in Chile (Figure 2). Similar results of the NA tree were observed for the internal genes (Supplementary figures 1-6), which suggested that A/swine/O’Higgins/VN1401-7826/2024 (H3N2) could correspond to a reassortant strain between seasonal human H3N2 and swine H1N2. It is crucial to mention all Chilean swine H1N2 strains present internal genes derived from the H1N1pdm09 lineage (9,11).

**Figure 1.**
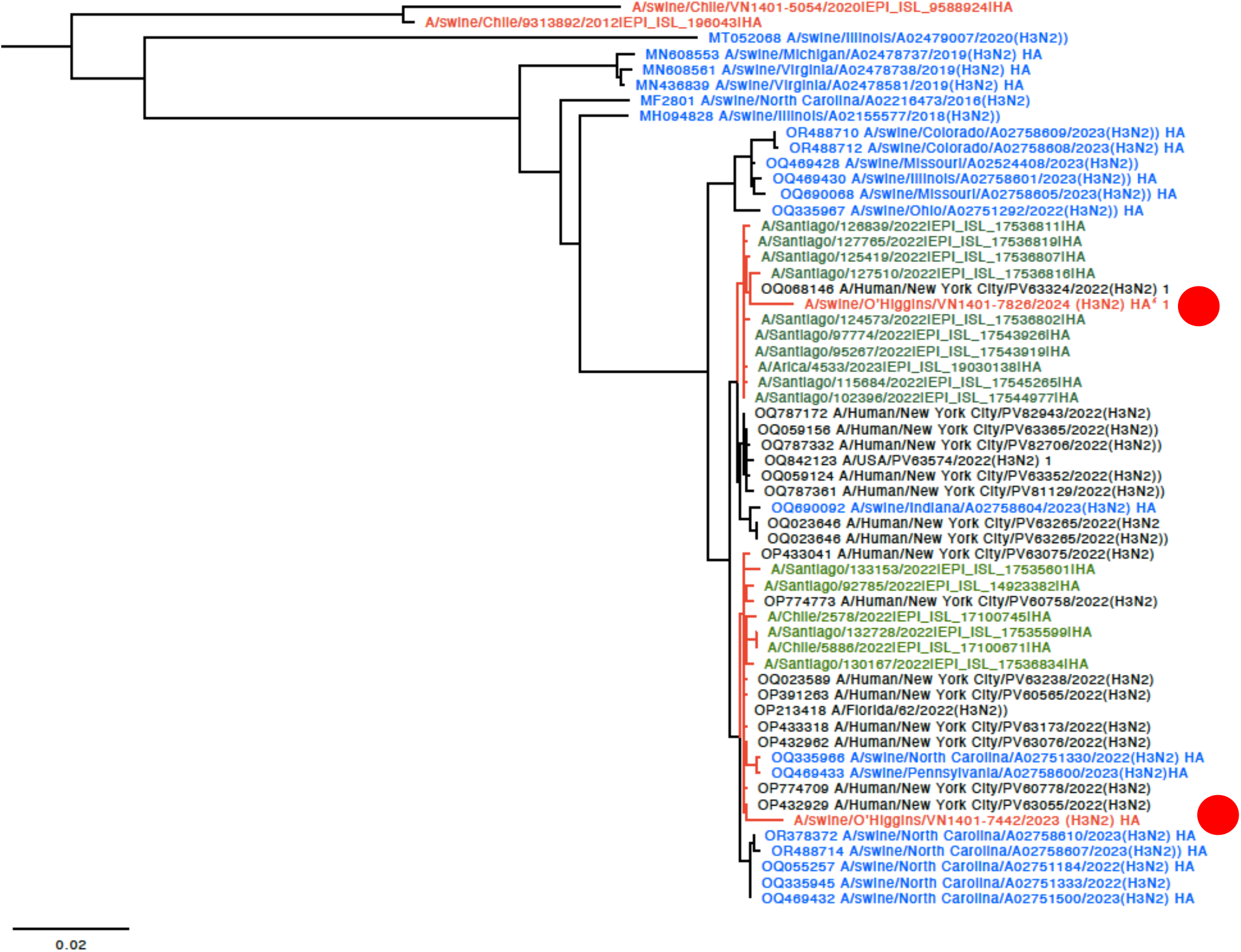
HA phylogenetic tree showcasing the introduction of two novel H3 viruses into Chile. The sequences are color-coded for clarity: red Chilean swine, green Chilean human, blue USA swine, and black indicates humans from the USA. For clarity, the novel Chilean swine isolates are depicted in a red circle.

**Figure 2.**
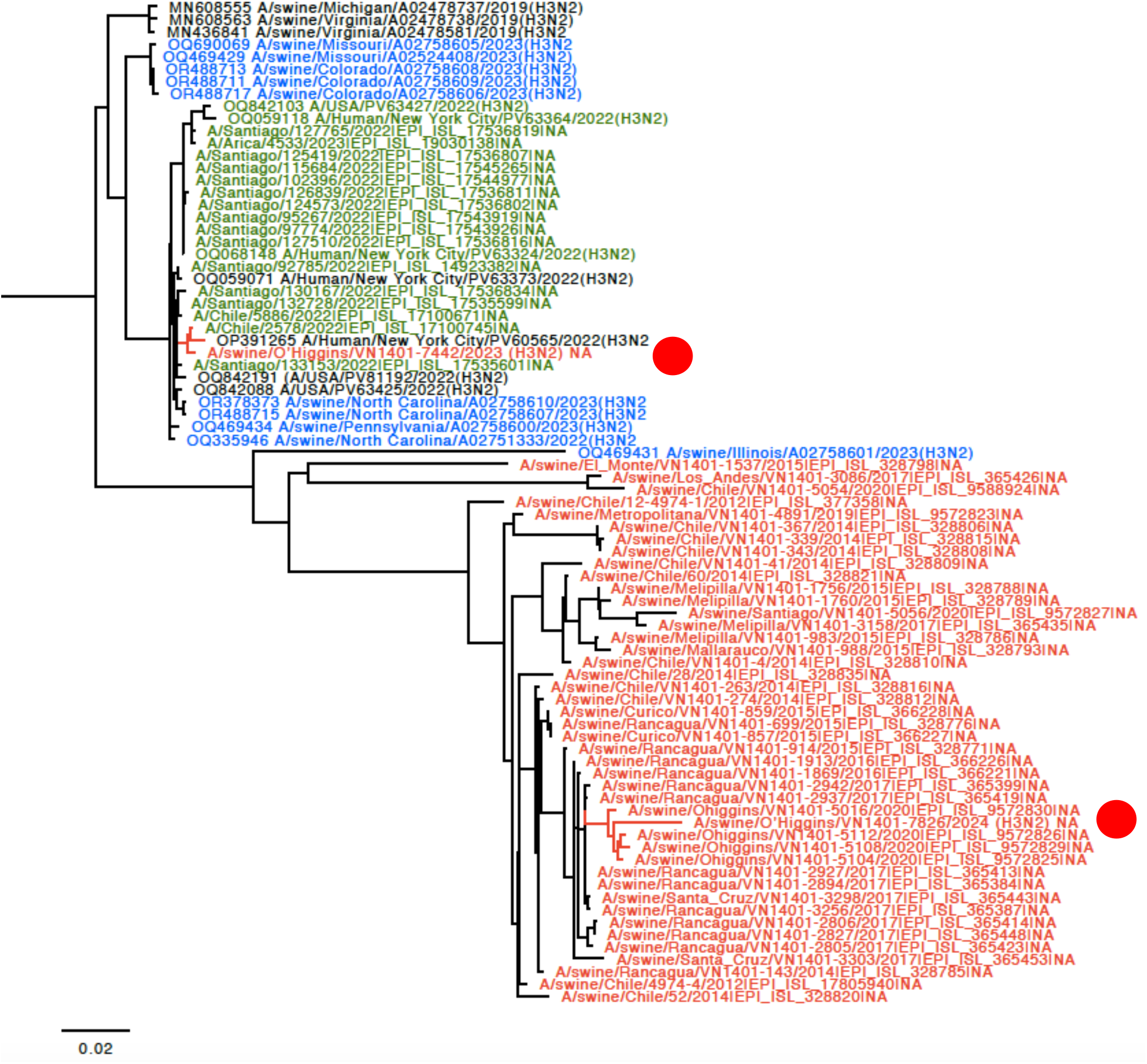
NA phylogenetic tree showcasing the introduction of N2 from human-origin novel viruses. The sequences are color-coded for clarity: red Chilean swine, green Chilean human, blue USA swine, and black indicates humans from the USA. For clarity, the novel Chilean swine isolates are depicted in a red circle.

Regarding reconfirming, if the A/swine/O’Higgins/VN1401-7826/2024 (H3N2) corresponded to a unique strain and no mixed infection, the assembling was repeated using a Chilean swine H1 reference sequence. We selected the A/swine/O’Higgins/VN1401-5112/2020 (H1N2) HA because the NA and internal genes of this strain are clustered with the 7826/2024 in all the segments (Figure 2, Supplementary figures 1-6). No contigs for H1 were obtained. The reassortment on the A/swine/O’Higgins/VN1401-7826/2024 (H3N2) was visualized by the recombination analysis of concatenated sequences identifying the A/Human/New York City/PV63324/2022(H3N2) as the minor parent (HA gene) and the A/swine/O’Higgins/VN1401-5112/2020 (H1N2) as the major parent (PB2, PB1, PA, NP, NA, M and NS) (Supplementary figure 7).

## 4 Discussion

In this study, we described the introduction of two H3N2 strains into the swine population in Chile. Notably, both strains have been associated with unexpected clinical outcomes in nursery farms endemic for the Influenza A virus, with both cases reported in 2023.

Phylogenetic analysis across all segments confirms that the strains are distinctly different. Furthermore, the absence of any known epidemiological link between them conclusively indicates that they are separate strains. However, genetic analysis reveals that both strains belong to the recent human seasonal H3 lineage. Specifically, the HA phylogeny strongly suggests these H3 strains were recently introduced by humans. The clustering of several human sequences from Chile with the novel swine H3 indicates a direct transmission from Chilean humans to Chilean swine farms (Figure 1). Reverse zoonotic is quite common in the Chilean swine population. Human-to-swine transmissions of the H1N1pdm09 strain have been extensively documented in Chile (8,9,11) and are also recognized globally (23). Other prevalent strains in Chilean swine include the H1N2s, which are similar to human strains. Previous research has shown that these H1N2 infections were the result of ancient human-to-swine transmissions. In contrast, to date, the H3N2 strain has been identified in only a few Chilean farms, which also corresponded to ancient introductions from humans (9).

For case 1, a complete genetic analysis of strain A/swine/O’Higgins/VN1401-7442/2023 (H3N2) confirms that all segments originated from human H3N2, confirming the spillover. For case 2, strain A/swine/O’Higgins/VN1401-7826/2024 (H3N2) shares the H3 human origin, but the rest of the genome includes a mix of internal genes from H1N1pdm09 and N2 from a prevalent H1N2, previously found in the same company. Despite these differences, the introduction of H3N2 into the swine population from humans remains a primary focus.

Over the past decade, there has been no clear dominance of any particular IAV subtypes in humans, with both H1N1pdm09 and seasonal H3N2 circulating (24), which alternates from year to year (25). For example, from August 2021 to December 2022 H3N2 predominated, while H1N1 became more prevalent globally, including in Chile, from March 2023 to January 2024 (GISAID Frequency Dashboard (v0.9)). Although there has been no evidence of novel H3N2 introductions into the human population for over a decade, H1N1 introductions have been frequently detected. Furthermore, the frequency of these subtypes in the human population does not necessarily correlate with their introduction into swine populations; however, requires more investigation.

Recent human-to-swine transmissions of the H3 strain have also been documented in the USA. Powell et al. (2024) reported several instances of the 2018-2019 human seasonal H3N2 influenza A virus spilling over into the swine population; however, these were not sustained within the pig populations (26). Subsequently, our research identified several seasonal human-like H3N2 viruses in the US pig population (OctoFLU), which closely resembled our findings. The closest strains from Indiana, Pennsylvania, and North Carolina clustered with the novel Chilean H3N2 viruses. These viruses appear not to be reassortant strains, similar to our Case 1, necessitating further studies to determine their persistence in the pig population. Conversely, from Case 2 the results suggest the presence of a real reassortant strain that was observed three months after the initial detection of H3 on the farm, strongly indicating that the virus has become established in the pig population. The sequenced sample corresponded to an isolate, therefore, there is a low possibility of reassortment during this process. Overall, both H3N2 introductions should be more studied in the future.

In both cases, there is a concern that the virus might spread to other farms within the same company through pig movements, warranting additional investigation. It is crucial to note, based on our historical data, that intra-species transmission of IAV in Chile is generally low among pig companies, with reverse zoonosis being of greater concern.

The presence of human-origin H3N2 viruses in the swine population might be more widespread globally than reported, with potential underreporting in certain regions due to limited surveillance or insufficient sequences being submitted to public databases. For instance, the swine IAV sequences available in South America to date total 459, while North America accounts for 19,287, representing 1.47% and 61.86% respectively of the total swine IAV viruses available on the GISAID repository. This significant discrepancy and low representativeness of this region extend and exacerbate in other areas with historic limitations on resources and research efforts, severely limiting our understanding of the transmission and evolution of influenza viruses across different geographical areas. Comprehensive sequencing requires specialized equipment, expertise, and resources; therefore, it is crucial to prioritize sequencing cases with unusual clinical presentations or characteristics that may indicate new viral introductions. As a surveillance enhancement alternative, Oxford Nanopore technologies have emerged, providing increased sequencing capabilities at a more economical cost and faster (14,27,28).

The Chilean government has strengthened its annual vaccination programs for personnel involved in swine and poultry operations. These efforts should be supplemented with additional biosecurity measures, including the use of masks and gloves, rapid testing for individuals suspected of being affected by influenza, and minimizing contact with pigs. These practices aim to reduce both reverse zoonotic and potential zoonotic events.

Despite comprehensive analysis of the whole genome and strong evidence indicating the introduction of seasonal human H3N2 into the Chilean pig population, this study has limitations. One significant limitation is that only one farm was followed closely and resample (case 2). Another area that could provide insights is time-lapse sequencing to determine the exact date of reassortment. Additionally, there was a failure to effectively subtype these new strains. Future research could address these gaps.

The implications of these findings extend beyond the swine industry, with potential implications for public health and disease control efforts. The results underscore the complex interplay between viral ecology, host specificity, and viral transmission dynamics. Understanding these dynamics is essential for assessing the risk of zoonotic transmission and developing effective strategies for disease prevention and control.

## 6 Author Contributions

Conceptualization, B.A. and V.N.; methodology, B.A., N.E., B.Q., F.B., and V.N.; formal analysis, B.A., N.A., and V.N.; writing—original draft preparation, B.A., and V.N.; writing—review and editing, V.N.; project administration, V.N..; funding acquisition, V.N., and R.M. All authors have read and agreed to the published version of the manuscript.

## 8 Acknowledgments

Thanks to Chilean swine veterinarians for sample collection. Thanks to Dr. Catalina Paro-Roa for her support in the sequencing process.

