## Supplemental for "Novel introductions of human-origin H3N2 Influenza viruses in Swine, Chile"

**Supplementary Table 1.-** Description of the total samples by type and their percentage of positivity to RT-PCR IAV.

| Sample | Total | RT-PCR IAV+ | Positivity |
|---|---|---|---|
| BS | 64 | 23 | 35,9% |
| L | 26 | 2 | 7,7% |
| NS | 1070 | 90 | 8,4% |
| OF | 32 | 12 | 37,5% |
| TS | 3 | 0 | 0,0% |
| **Totals** | **1195** | **127** | **10,6%** |

**Supplementary Table 2.-** Description of the closest sequences detected with BLASTn for each genome segment of the A/swine/O'Higgins/VN1401-7442/2023 and A/swine/O'Higgins/VN1401-7826/2024 viruses.

| **Sequence BLASTn** | **A/swine/O'Higgins/VN1401-7442/2023** | | | | **A/swine/O'Higgins/VN1401-7826/2024** | | | |
|---|---|---|---|---|---|---|---|---|
| **Segment** | **Strain name** | **Accesion number** | **Host** | **Percent identity** | **Strain name** | **Accesion number** | **Host** | **Percent identity** |
| **HA** | A/Human/New York City/PV63055/2022(H3N2)) segment 4 hemagglutinin (HA) gene, complete cds. | OP432929.1 | Human | 99.26% | (A/Human/New York City/PV63324/2022(H3N2)) segment 4 hemagglutinin (HA) gene, complete cds. | OQ068146 | Human | 98.98% |
| **NA** | Influenza A virus genome assembly, segment: 6. | OY283278 | Human | 99.66% | A/swine/Ohiggins/VN1401-5105/2020(HxN2)) segment 6 neuraminidase (NA) gene, complete cds. | MZ005159 | Swine | 97.20% |
| **PB2** | Influenza A virus (A/Human/New York City/PV63055/2022(H3N2)) segment 1 polymerase PB2 (PB2) gene, complete cds | OP432930.1 | Human | 99.83% | Influenza A virus (A/swine/Ohiggins/VN1401-5016/2020(H1N2)) segment 1 polymerase PB2 (PB2) gene, complete cds | MZ005153.1 | Swine | 97.52% |
| **PB1** | A/Human/New York City/PV63357/2022(H3N2)) segment 2 polymerase PB1 (PB1) and PB1-F2 protein (PB1-F2) genes, complete cds. | OQ059059 | Human | 99.91% | A/swine/Ohiggins/VN1401-5108/2020(H1N2)) segment 2 polymerase PB1 (PB1) gene, complete cds; and nonfunctional PB1-F2 protein (PB1-F2) gene, complete sequence. | MZ013916 | Swine | 98.12% |
| **PA** | Influenza A virus (A/Human/New York City/PV82899/2022(H3N2)) segment 3 polymerase PA | OQ787306.1 | Human | 99.69% | Influenza A virus (A/swine/Ohiggins/VN1401-5101/2020(H1N1)) segment 3 polymerase PA (PA) and | MZ005177 | Swine | 97.94% |

| | | | | | | | | |
|---|---|---|---|---|---|---|---|---|
| | (PA) and PA-X protein (PA-X) genes, complete cds | | | | PA-X protein (PA-X) genes, complete cds. | | | |
| **NP** | Influenza A virus (A/Human/New York City/PV63367/2022(H3N2)) segment 5 nucleocapsid protein (NP) gene, complete cds | OQ059172.1 | Human | 99.55% | A/swine/Ohiggins/VN1401-5092/2020(HxN2)) segment 5 nucleocapsid protein (NP) gene, complete cds. | MZ005183 | Swine | 98.34% |
| **M** | Influenza A virus genome assembly, segment: 7. | OX411443.1 | Human | 99.90% | Influenza A virus (A/swine/Ohiggins/VN1401-5108/2020(H1N2)) segment 7 matrix protein 2 (M2) and matrix protein 1 (M1) genes, complete cds | MZ005134.1 | Swine | 98.83% |
| **NS** | A/WA/31872/2022(H3N2)) segment 8 nuclear export protein (NEP) and nonstructural protein 1 (NS1) genes, complete cds. | OQ180177.1 | Human | 99.66% | Influenza A virus (A/swine/Rancagua/VN1401-2807/2017(H1N2)) segment 8 nuclear export protein (NEP) and nonstructural protein 1 (NS1) genes, complete cds | MH346870.1 | Swine | 98.54% |

Figure 1. PB2 phylogenetic tree. The sequences are color-coded for clarity: red Chilean swine, green Chilean human, blue USA swine, and black indicates humans from the USA. For clarity, the isolate A/swine/O'Higgins/VN1401-7826/2024 (H3N2) is depicted with a red circle.

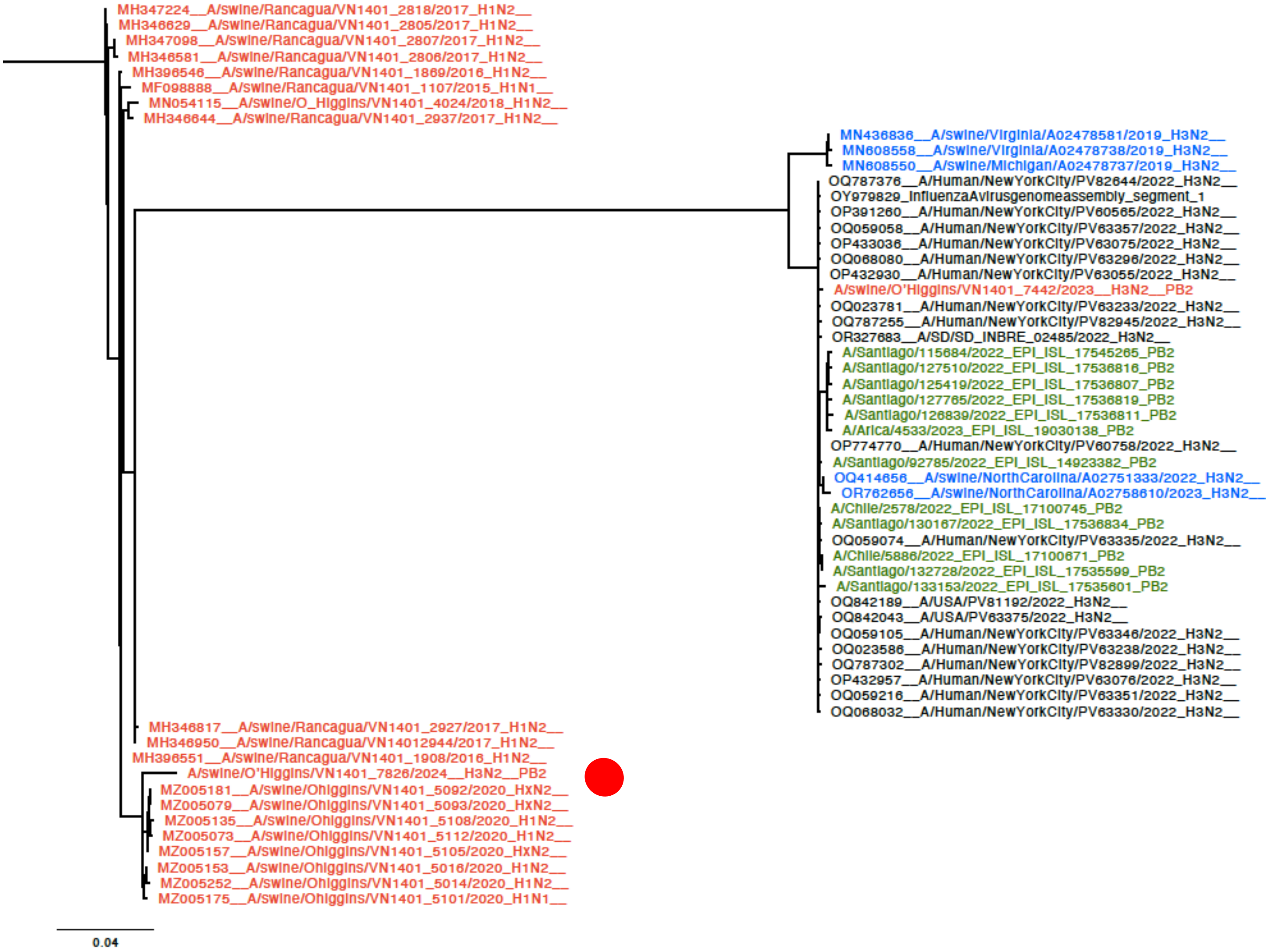

Figure 2. PB1 phylogenetic tree. The sequences are color-coded for clarity: red Chilean swine, green Chilean human, blue USA swine, and black indicates humans from the USA. For clarity, the isolate A/swine/O'Higgins/VN1401-7826/2024 (H3N2) is depicted with a red circle.

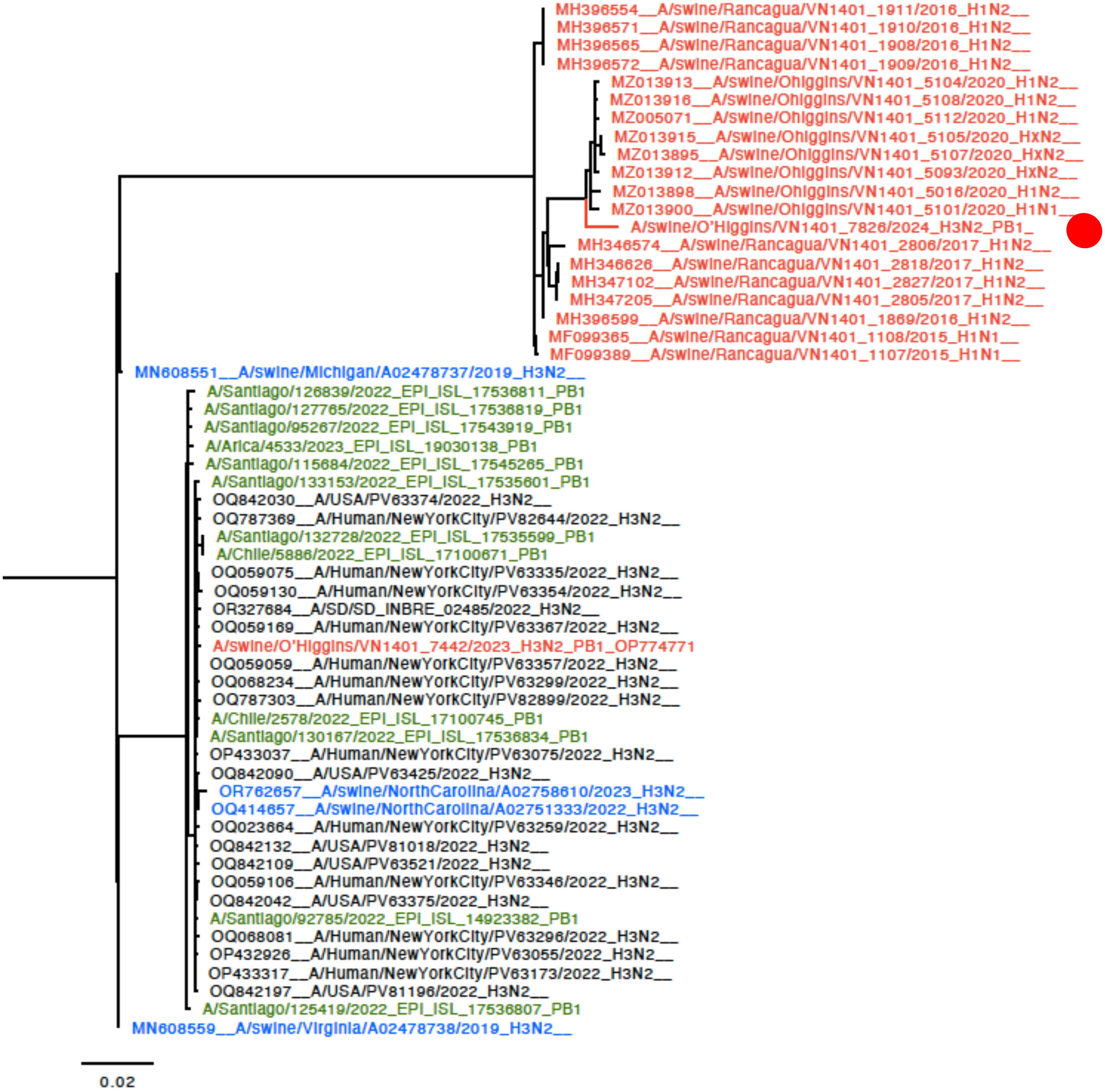

Figure 3. PA phylogenetic tree. The sequences are color-coded for clarity: red Chilean swine, green Chilean human, blue USA swine, and black indicates humans from the USA. For clarity, the isolate A/swine/O'Higgins/VN1401-7826/2024 (H3N2) is depicted with a red circle.

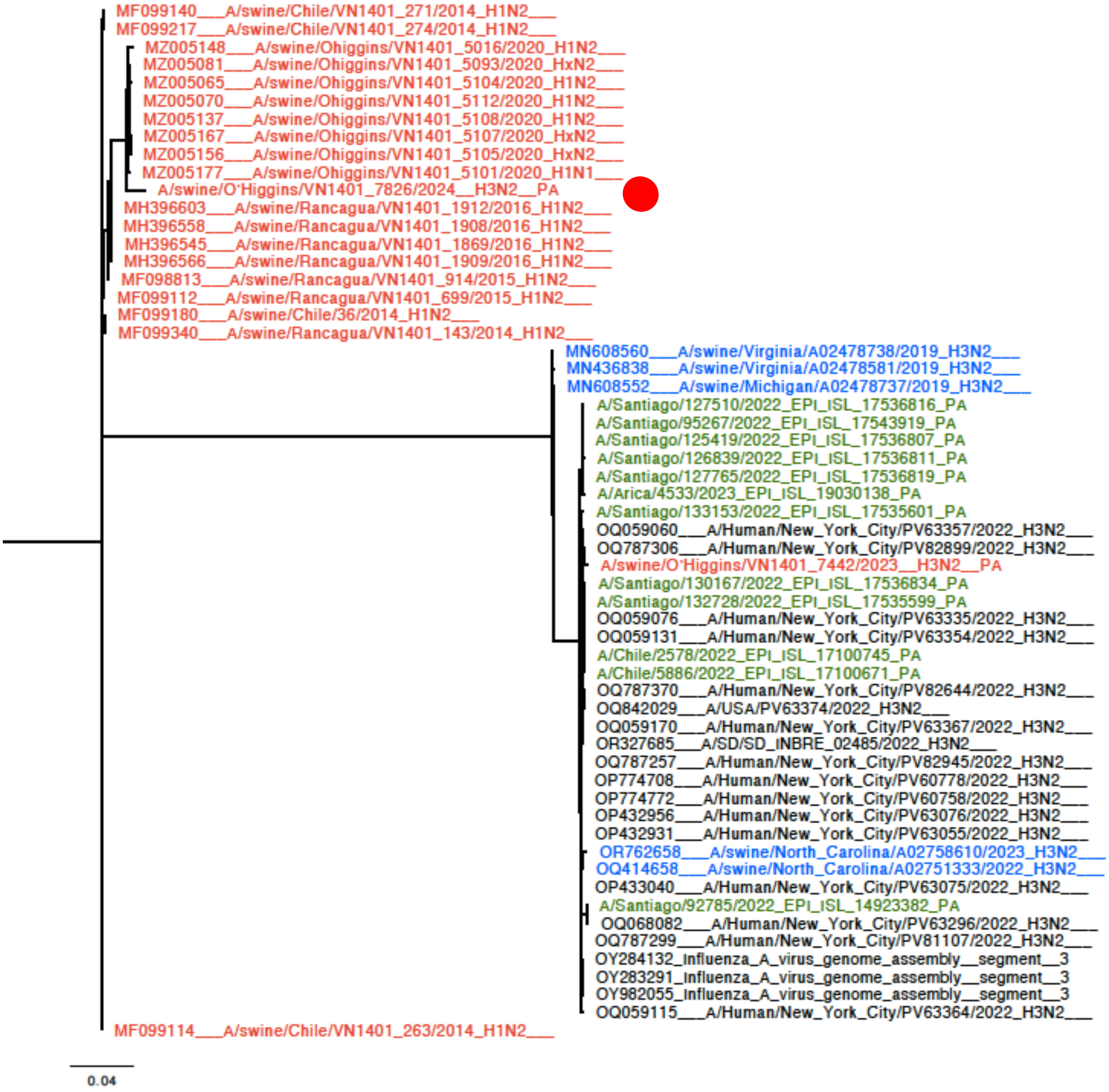

Figure 4. NP phylogenetic tree. The sequences are color-coded for clarity: red Chilean swine, green Chilean human, blue USA swine, and black indicates humans from the USA. For clarity, the isolate A/swine/O'Higgins/VN1401-7826/2024 (H3N2) is depicted with a red circle.

Figure 5. M phylogenetic tree. The sequences are color-coded for clarity: red Chilean swine, green Chilean human, blue USA swine, and black indicates humans from the USA. For clarity, the isolate A/swine/O'Higgins/VN1401-7826/2024 (H3N2) is depicted with a red circle.

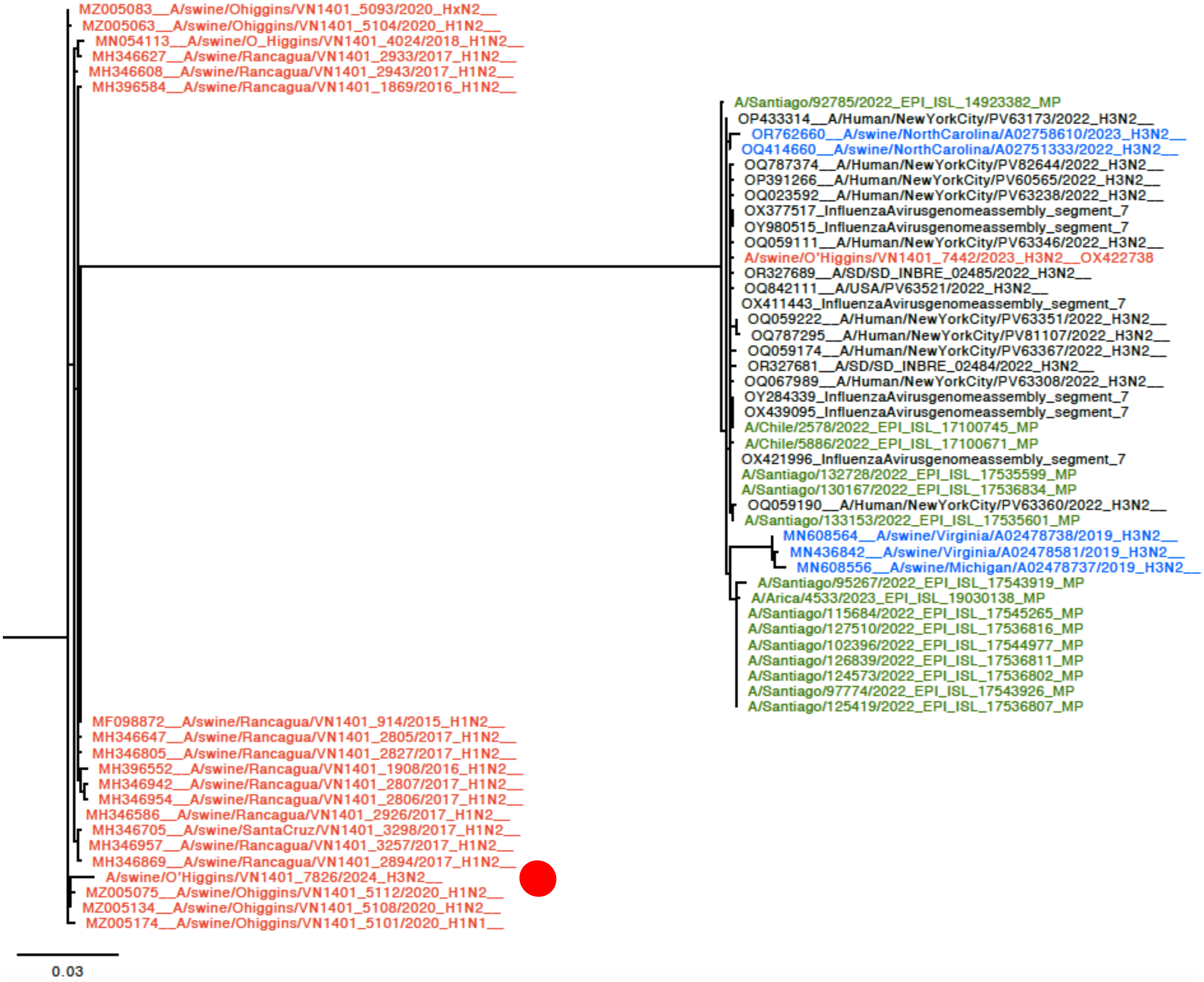

Figure 6. NS phylogenetic tree. The sequences are color-coded for clarity: red Chilean swine, green Chilean human, blue USA swine, and black indicates humans from the USA. For clarity, the isolate A/swine/O'Higgins/VN1401-7826/2024 (H3N2) is depicted with a red circle.

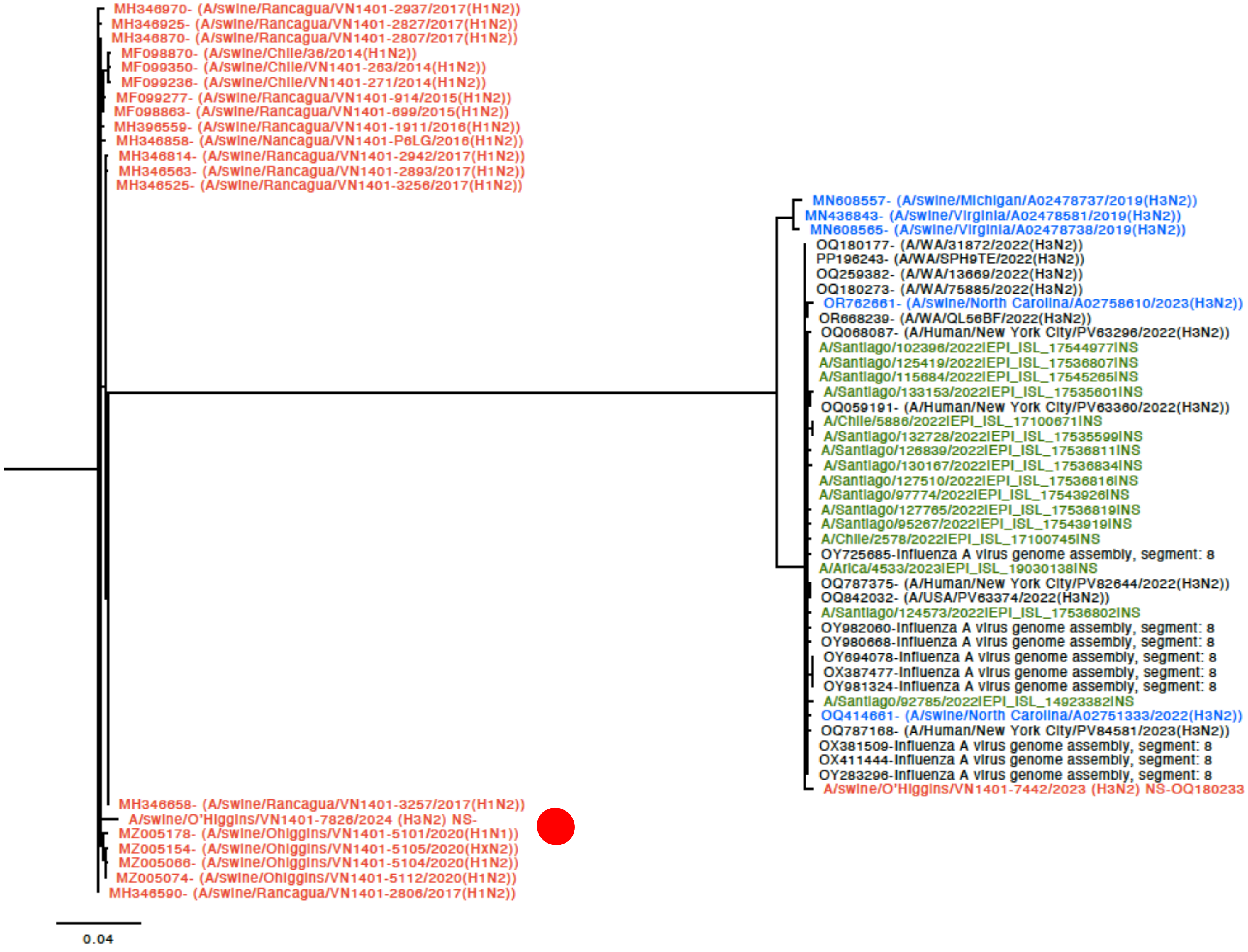

Figure 7. Dual Brothers recombination analysis evidence that A/swine/O’Higgins/VN1401-7826/2024 (H3N2) is a reassortant strain in the HA gene.

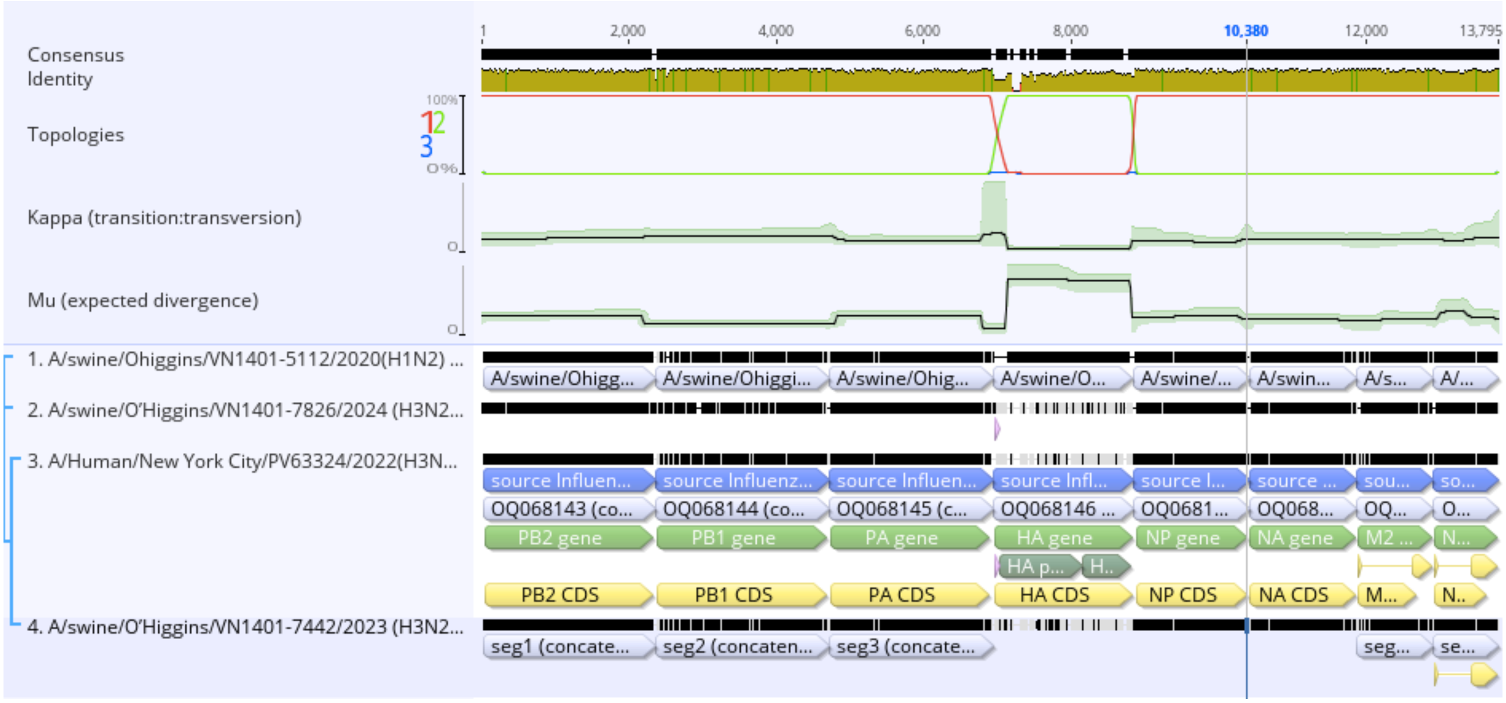
